# Parallel Evolution of Cold Tolerance Within *Drosophila melanogaster*

**DOI:** 10.1101/063545

**Authors:** John E. Pool, Dylan T. Braun, Justin B. Lack

## Abstract

*Drosophila melanogaster* originated in tropical Africa before expanding into strikingly different temperate climates in Eurasia and beyond. Here, we show that elevated cold tolerance has arisen at least three times within this species: beyond the well-studied non-African case, we show that populations from the highlands of Ethiopia and South Africa have significantly increased cold tolerance as well. We observe greater cold tolerance in outbred versus inbred flies, but only in populations with higher inversion frequencies. Each cold-adapted population shows lower inversion frequencies than a closely-related warm-adapted population, suggesting that inversion frequencies may decrease with altitude in addition to latitude. Using the *F_ST_*-based “Population Branch Excess” statistic (*PBE*), we found only limited evidence for parallel genetic differentiation at the scale of ~4 kb windows, specifically between Ethiopian and South African cold-adapted populations. And yet, when we looked for single nucleotide polymorphisms (SNPs) with codirectional frequency change in two or three cold-adapted populations, strong genomic enrichments were observed from all comparisons. These findings could reflect an important role for selection on standing genetic variation leading to “soft sweeps”. One SNP showed sufficient codirectional frequency change in all cold-adapted populations to achieve experiment-wide significance: an intronic variant in the synaptic gene *Prosap*. More generally, proteins involved in neurotransmission were enriched as potential targets of parallel adaptation. The ability to study cold tolerance evolution in a parallel framework will enhance this classic study system for climate adaptation.

## INTRODUCTION

Thermal tolerance is a critical determinant of species ranges and organismal adaptation to environments. Adaptation to cold climates and its genetic basis has been a topic of particular scientific interest, including studies of humans (*e.g.* Hancock *et al.* 2010) and *Drosophila* (*e.g.* Hoffmann *et al.* 2003; Ayrinhac *et al.* 2004; Svetec *et al.* 2011). Unraveling the genetic architecture and biological mechanisms underlying insect cold tolerance may be especially relevant for identifying factors that currently constrain the geographic distributions of medically important insects such as disease vectors.

Originating from a relatively warm sub-Saharan ancestral range (Lachaise *et al.* 1988; Pool *et al.* 2012), *D. melanogaster* ultimately became associated with human settlements and spread throughout the world, including environments much colder than its original habitat. Evidence for this species’ genetic response to novel thermal environments includes geographic variation in cold tolerance (Hoffmann *et al.* 2003) and the presence of parallel allele frequency clines in North America and Australia (Turner *et al.* 2009). These clines provide striking examples of rapid evolution, in that North American and Australian populations of *D. melanogaster* appear to have descended from the admixture of European and African source populations within the past century or two (Keller 2007; Bergland *et al.* 2016). Although source population ancestry shows a genome-wide latitude cline on these continents (Kao *et al.* 2015; Bergland *et al.* 2016), loci showing extreme latitude clines could reflect the locally adaptive reassortment of pre-existing genetic variants that differed between the African and European source populations.

In contrast, the earlier expansion of *D. melanogaster* from Africa into Eurasia, which occurred roughly 10,000 years ago (Thornton and Andolfatto 2006; Duchen *et al.* 2013), may have involved novel genetic adaptation to cold environments. This adaptation is most commonly studied via comparisons between European and sub-Saharan African populations (*e.g.* Kauer *et al.* 2002; Svetec *et al.* 2011; Božičević *et al.* 2016). However, the species’ sub-Saharan exodus entailed a strong population bottleneck that complicates the population genetic detection of adaptive differences between sub-Saharan and European populations, by stochastically increasing the level of neutral genome-wide differentiation between these populations. We suggest that the genetic basis of European cold adaptation could alternatively be studied by comparison with a population that had gone through the same bottleneck, but showed less adaptation to cold climates. Northern African populations appear to be genetically similar to European populations (Dieringer *et al.* 2005; Lack *et al.* 2015), and in this study we compare fly strains from France and Egypt.

Although populations from outside sub-Saharan Africa are known to offer a range of opportunities for studying cold tolerance evolution, these populations may largely reflect a single origin of cold tolerance. In contrast, sub-Saharan Africa could offer previously unappreciated independent origins of cold adaptation. Africa’s most extensive highlands are found in Ethiopia, featuring a consistently cool environment. Seasonally cold environments are found in South Africa due to a combination of higher altitude and latitude. We therefore study low and high altitude pairs of populations from each of these countries, representing warmer and cooler environments respectively (Figure 1).

**Figure 1.**
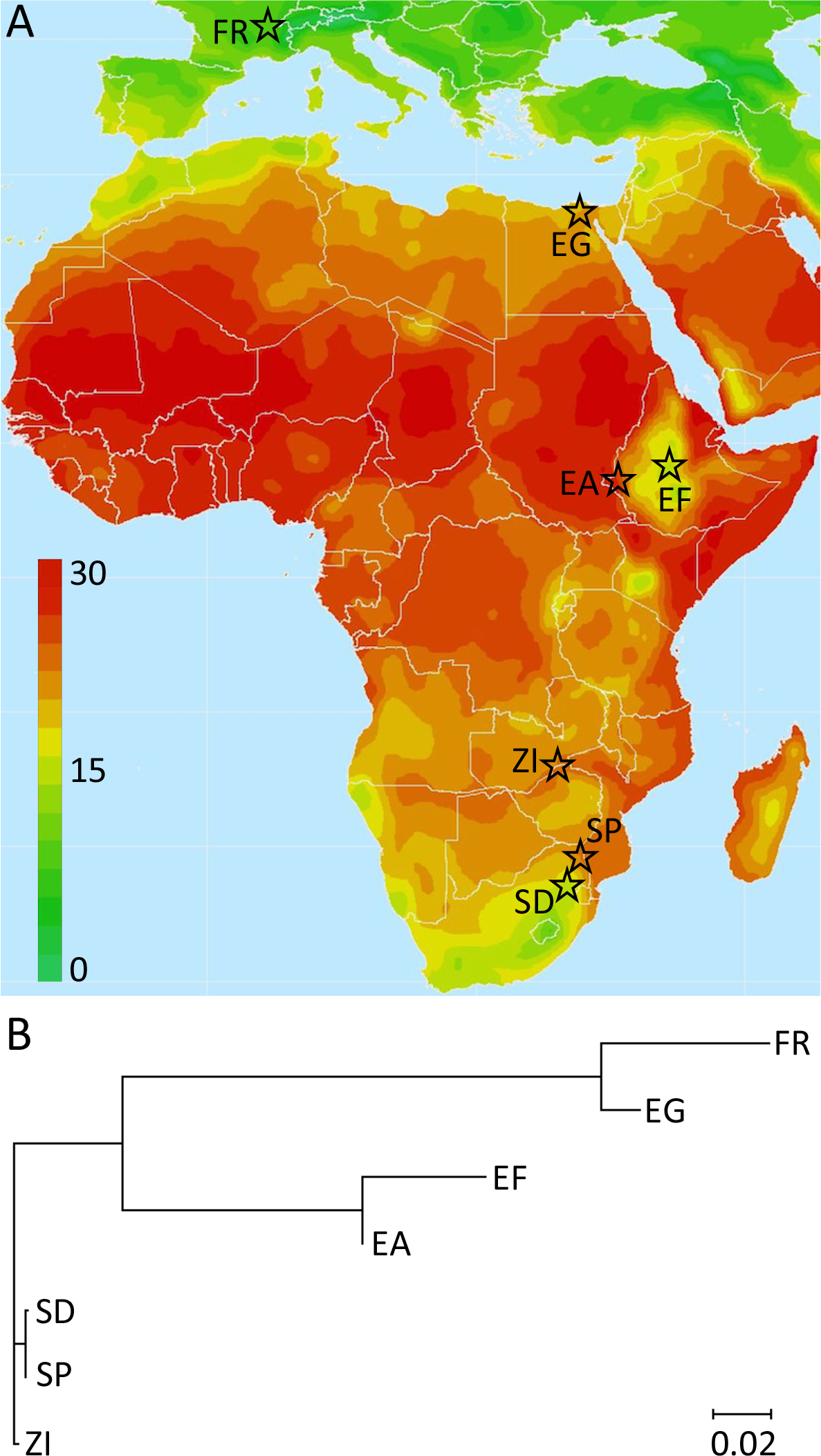
(A) Geographic locations and thermal environments of the studied populations are depicted. We compare three pairs of cold-/warm-adapted populations: France/Egypt and highland/lowland populations from Ethiopia and from South Africa. The origin of the ancestral range Zambia sample used in population genomic analyses is shown as well. The source map indicating average temperature (°C) is courtesy of The Nelson Institute, Center for Sustainability and the Global Environment, UW-Madison. (B) A neighbor-joining tree, based on transformed genome-wide average *F_ST_* values (Cavalli-Sforza 1969) and rooted along the branch to ZI, was generated using MEGA 7.0.16 (Kumar *et al* 2016).

If confirmed, the existence of cold-tolerant populations in each of these three regions would appear to indicate parallel phenotypic evolution – that is, each cold-adapted population being founded independently by warm-adapted progenitors. *D. melanogaster* is thought to originate from the southern portion of Africa (Pool *et al.* 2012), and thus cold-adapted populations from this region could be the most ancient. However, the species’ northward expansion within Africa would have originated far from South Africa. Instead, populations would have expanded from tropical environments to ultimately occupy Ethiopia and Eurasia. Population genomic data suggest that the species’ colonization of Ethiopia and Eurasia involved separate expansions, in that Ethiopian populations do not appear to be the source of the out-of-Africa emigration (Pool *et al.* 2012).

Parallel phenotypic evolution can be explained by at least three distinct genetic scenarios. First, cold tolerance could evolve from independent mutations in each population, at either the same or different genes. Second, cold tolerance could arise from a common pool of standing genetic variation present in the ancestral population. Third, cold tolerance variants could be transmitted from one population to another by migration. Although we argue that cold tolerance was favored independently in each of these regions, gene flow has occurred (Pool *et al.* 2012) and could contribute to cold tolerance evolution.

Populations in Europe and in the highlands of Ethiopia and South Africa have each experienced distinct selective pressures related to their abiotic and biotic environments. However, the component of each population’s local adaptation that relates to cold tolerance (or correlated selective pressures) could produce a genomic signal of genetically parallel evolution, if the same genes tend to contribute in each population. After documenting cold tolerance evolution in these three regions, we then tested for parallel genetic differentiation specific to the cold-adapted populations. Limited evidence of parallel genetic differentiation was obtained from window-based analysis of allele frequency change. However, a strong genomic enrichment of SNPs with codirectional allele frequency change was observed between all cold-adapted populations, including SNPs in genes including membrane-bound and neurotransmission-related genes.

## RESULTS

### Cold tolerance has evolved in three geographic regions

We used three pairs of population samples to search for evidence of the parallel phenotypic evolution of cold tolerance within *D. melanogaster* (Figure 1). Our assay consisted of the proportion of female flies still standing after 96 hours at 4°C. Results indicated highly significant increases in cold tolerance for strains originating from colder environments in France, Ethiopia, and South Africa (Table 1; Table S1). The France strains showed the greatest degree of cold tolerance. This population represents the adaptation of *D. melanogaster* to temperate Eurasian climates, which has been a rich source of biological insight. Cold-tolerant European *D. melanogaster* appear to have made substantial genetic contributions to recently founded populations from the Americas and Australia as well (Nunes *et al.* 2008; Bergland *et al.* 2016). Importantly, our Egypt sample lacks most of the cold tolerance present in France, in spite of these samples’ low genetic differentiation (Lack *et al.* 2015). This pair of European and northern African populations therefore represents an excellent comparison for studies of cold adaptation.

**Table 1.**
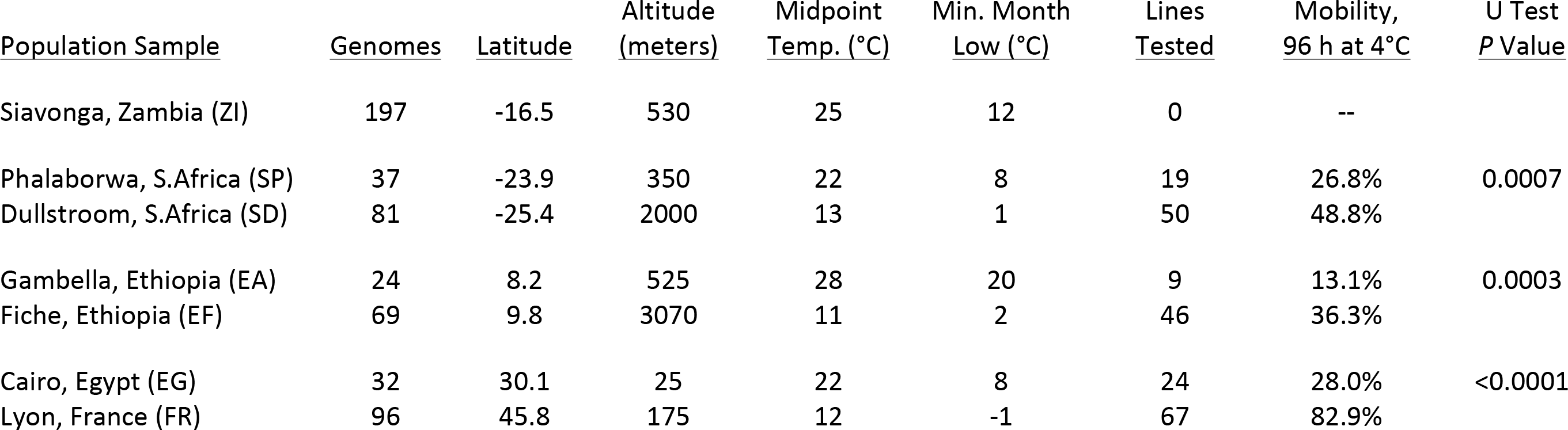
*D. melanogaster* inbred lines from three distinct regions show elevated cold tolerance (measured as the percentage of female flies still standing or moving after 96 h at 4°C). To illustrate climate differences, estimates of the “midpoint temperature” (the average of 24 monthly average highs and lows) and the minimum monthly average low are given. *P* values show that within each pair of populations, the sample originating from the higher altitude/latitude shows significantly greater cold tolerance.

The increased cold tolerance detected in Ethiopian and South African populations (Table 1) has not been previously studied, and is likely to reflect independent natural selection in each case (Introduction). The degree of cold tolerance in these populations is less than that observed for France, which might be explained by the presence of seasonally much colder environments in parts of Europe, and perhaps by the smaller geographic size of cool regions in Africa (in light of the balance between local adaptation and gene flow). Still, the Ethiopian and South African population comparisons are both strongly significant, with greater than 20% absolute increases in the fraction of flies standing after 96 hours at 4°C.

### Variable Influence of Inbreeding on Cold Tolerance

In a limited number of cases, we also investigated the cold tolerance of outbred flies from each population. For these within-population crosses, males from different inbred strains were crossed to females from a single reference strain. Results were surprisingly population-dependent: compared with expectations based on the average of the parental inbred strains, three populations had significantly improved cold tolerance after outbreeding (Egypt and the two South African samples), while the other three populations showed no trend in this direction (Table 2; Table S2). Although the numbers of crosses are modest, this can not account for the divergent pattern observed; the null probability for at least three populations to have a significant increase in outbred tolerance (*P* < 0.05), with the rest showing no trend (*P* ≥ 0.5) is quite low (*P* = 0.0026, based on one million randomizations of the direction of change between outbred and midparent tolerance values). Hence, the influence of outbreeding on cold tolerance vary considerably depending on the population (or based on the specific reference strain used in our crosses). Thus, the effects of inbreeding on tolerance traits may depend not only on the trait examined (Dahlgaard and Hoffmann 2000), but also on the genotypes being studied. One potential factor in our population-sensitive outbreeding results is inversion frequencies: the three populations with increased cold tolerance after outbreeding were the also the three populations with the highest frequency of inverted chromosomes (see below). This tentative correlation might be explained by associative overdominance in connection with recessive deleterious alleles carried on inverted chromosomes (Ohta 1971).

**Table 2.** The relationship between inbred and outbred cold tolerance differs strongly population. Here, outbred cold tolerance (measured as the percentage of female flies s standing or moving after 96 h at 4°C) is compared with the midparent values expected based on the average cold tolerance of the parental inbred strains. *P* values indicate st advantages of outbred flies for EG, SD, and SP, but no hint of such an effect for the othe populations. These same three populations have the highest inversion frequencies (Fig 2).

| <u>Population</u> | <u>Midparent</u><br><u>Mobility</u> | <u>Outbred</u><br><u>Mobility</u> | <u>Number</u><br><u>of Crosses</u> | <u>Outbred</u><br><u>Greater</u> | <u>Binomial</u><br><u>P Value</u> |
| --- | --- | --- | --- | --- | --- |
| S. Africa SP | 11.8% | 50.1% | 6 | 6 | 0.016 |
| S. Africa SD | 51.5% | 73.0% | 6 | 6 | 0.016 |
| Ethiopia EA | 12.8% | 11.8% | 4 | 2 | 0.688 |
| Ethiopia EF | 53.4% | 50.6% | 12 | 4 | 0.927 |
| Egypt EG | 28.3% | 41.8% | 11 | 9 | 0.033 |
| France FR | 91.4% | 92.4% | 7 | 4 | 0.500 |

### Parallel inversion frequency change in cold-adapted populations

We showed above that populations in Ethiopia, France, and South Africa have experienced parallel phenotypic evolution toward greater cold tolerance. When then sought to address two questions regarding the evolutionary genetic mechanisms underlying parallel cold tolerance evolution. First, is there evidence that the same genes have contributed to adaptive evolution in two or more cold-adapted populations? And is there evidence that the same variants have been independently selected?

Before proceeding to full scans of genomic variation, we assessed whether inversion polymorphisms showed evidence for parallel frequency change in cold-adapted populations. Based on PCR-testing for eight polymorphic inversions, we found a trend toward lower inversion frequencies in all three cold-adapted populations (Table S3). In particular, the five most common inversions all showed lower frequency in at least two cold-adapted populations (Figure 2). Overall, fewer chromosome arms carried inversions in SD relative to SP (32% vs. 37%, *P* = 0.26), EF relative to EA (6% vs. 17%, *P* = 0.005), and FR relative to EG (16% vs. 47%, *P* < 0.0001). In order to separate inversion-linked frequency changes from the potential influence of selection on specific genes and variants, we therefore excluded all genomes carrying an inversion on a given chromosome arm from the analyses described below. These findings are concordant with past results on latitudinal inversion clines (*e.g*. Lemeunier and Aulard 1992; Kapun *et al.* 2016), and provide evidence for independent decreases in inversion frequency in cool high altitude regions.

**Figure 2.**
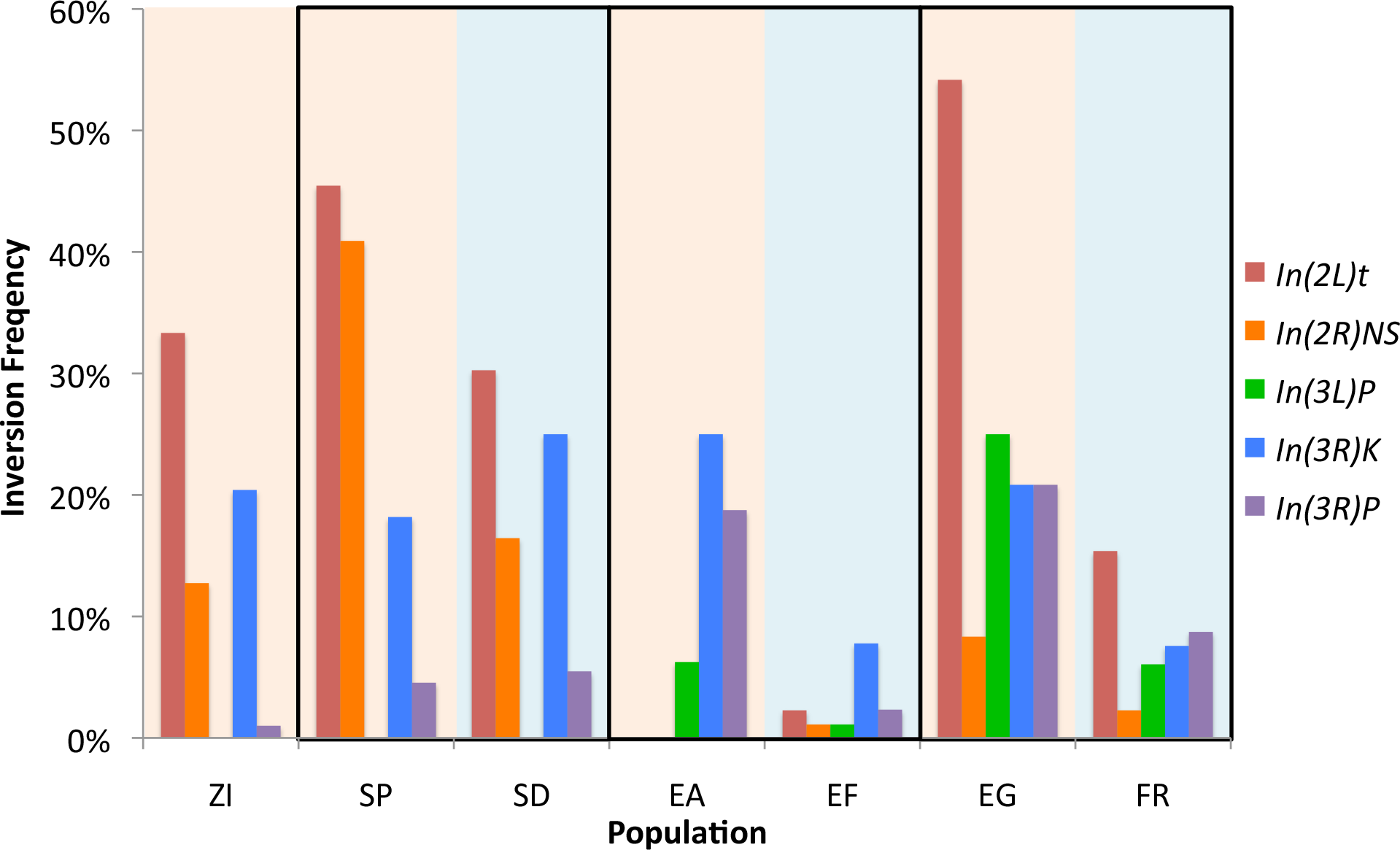
The frequencies of the five most common inversion polymorphisms tend to decrease in cold-adapted populations (blue background) relative to closely-related warm-adapted populations (tan background in same box). Further results of this inversion PCR testing are given in Table S3.

### Genomic window analysis of parallel genetic differentiation

The population genetic signals of adaptive genetic differentiation can be variable. Selective sweeps may be complete or incomplete. Further, the outcome of positive selection may be described as either a hard sweep (when a single haplotype rises in frequency, due to a new mutation or potentially a rare variant) or as a soft sweep (when multiple haplotypes rise in frequency, due to standing genetic variation or recurrent mutation). When statistics comparing genetic variation between two populations are applied to chromosomal windows, sensitivity is highest for complete hard sweeps; and depending on the parameters, significant power may persist for incomplete and/or soft sweeps as well (Lange and Pool 2016). However, soft sweeps involving many unique haplotypes may lead to high genetic differentiation at the causative SNP alone (and perhaps a handful of linked variants), a pattern which may be undetectable by window scans.

We therefore pursued both window and single SNP analyses of parallel genetic differentiation. For both analyses, we used a recently described *F_ST_*-based statistic known as *Population Branch Excess* (*PBE*; Yassin *et al.* 2016). Like the related *PBS* (Yi *et al.* 2010), *PBE* uses *F_ST_* values among three populations. Whereas *PBS* simply quantifies the length of the focal population’s branch on a genetic distance tree, *PBE* asks whether this *PBS* value is unexpectedly high in light of (1) *F_ST_* between the two non-focal populations at this locus, and (2) patterns of genetic differentiation across other loci (Materials and Methods). The most relevant difference between *PBE* and *PBS* is their behavior at loci that are subject to high genetic differentiation between all pairs of populations (due to recurrent positive selection not specific to the focal population, or other forces such as background selection). *PBS* should detect such loci as outliers for high genetic differentiation, but *PBE* is designed not to. Instead, *PBE* responds most strongly to cases where genetic differentiation is elevated only between the focal population and each of the others. *PBE* fits our interest in detecting loci that are specifically under selection in the cold-adapted populations, while avoiding false signals of genetically parallel cold adaptation from loci with globally high *F_ST_*.

We first evaluated *PBE* in ~4 kb windows, separately for each population pair, with the ancestral range Zambia sample as the third population in each case. Note that selection in either of the non-focal populations generates negative *PBE* values, rather than positive outliers. We then tested for a significant overlap in (positive) *PBE* outliers between different cold-adapted populations, using a permutation approach. No significant overlap in window *PBE* outliers was detected when France was compared against Ethiopia (*P* = 0.36) or South Africa (*P* = 0.94). But when the two cold-adapted sub-Saharan populations were compared, a significant overlap in *PBE* outliers was detected (*P* = 0.013). The latter result reflected a moderate enrichment of 77 overlapping outlier regions compared to a random expectation of 61. Thus, window *PBE* provided some evidence of parallel adaptation, but only between the two sub-Saharan population pairs.

### Genomic variant analysis of parallel genetic differentiation

Particularly if natural selection has acted on standing genetic variation, it is possible that individual SNPs might show parallel frequency changes without strong effects on linked variants. We therefore evaluated site-specific *PBE* and tested whether SNPs with strong cold vs. warm frequency differences in two different population pairs tended to be “codirectional” (the same allele increased in both cold-adapted populations, relative to each one’s warm-adapted counterpart) rather than “antidirectional”. In contrast to the window analysis described above, all three comparisons of population pairs showed a genomic signal of parallel SNP frequency change (Figure 3), in that SNPs with high *PBE* quantiles in two different population pairs were more likely to involve codirectional frequency change. Surprisingly, the strongest enrichment of codirectional *PBE* SNP outliers did not come from Ethiopia and South Africa (the comparison that had significantly overlapping window *PBE* outliers). Instead, Ethiopia and France/Egypt showed the greatest excess of codirectional vs. antidirectional SNPs that had *PBE* quantile below 0.05 in both pairs (Figure 3). Stronger enrichments were observed for more stringent *PBE* thresholds: for example, the numbers of codirectional versus antidirectional SNPs with *max*(*PBEQ*) < 0.005 was 45 vs. 8 for SD/EF, 7 vs. 1 for SD/FR, and 21 vs. 2 for EF/FR (Table S4). As expected based on linkage, some codirectional outliers were clustered, but overall they were widely distributed across the analyzed genomic regions (Figure S1, Table S5).

**Figure 3.**
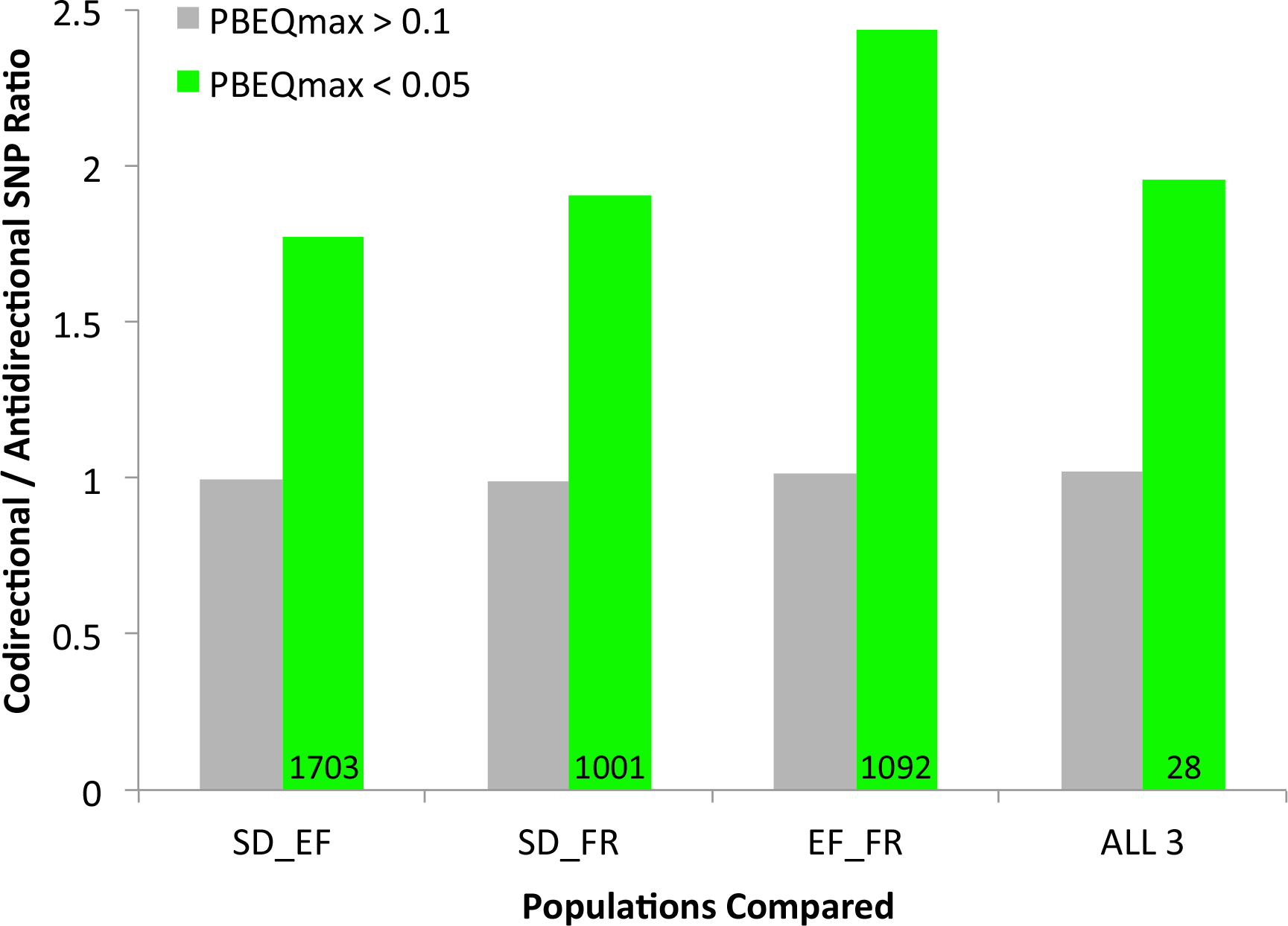
The genomic enrichment of SNPs with parallel frequency differences between cold-adapted and warm-adapted populations is illustrated. A SNP is classified as codirectional if the same allele is at higher frequency in cold-adapted populations compared to their warm-adapted counterparts. The ratio of codirectional to antidirectional SNPs is depicted for two classes of sites. Green bars indicate variants with high *PBE* values in each population pair considered (SNP *PBE* quantile below 0.05), and the total number of such codirectional SNPs is shown. Gray bars are for SNPs with *PBE* quantile above 0.1 for at least one of the population pairs. Joint *PBE* outliers consistently show an excess of codirectional SNPs, while non-outliers show very similar numbers of codirectional and antidirectional SNPs. Note that for the comparison including all three warm and cold populations pairs (right), ratios were multiplied by three to account for the expectation that only one quarter of SNPs should be codirectional by chance.

If the above patterns were produced by elevated gene flow between warm-adapted populations or between cold-adapted populations, then a general excess of codirectional over antidirectional SNPs would be expected. In contrast to this prediction, SNPs that were not outliers in both population pairs were equally likely to be codirectional or antidirectional in all comparisons (Figure 3). While the stochastic variance induced by population bottlenecks might modestly increase the number of codirectional outliers (Tennessen and Akey 2011), genetic differentiation between the cold- and warm-adapted populations of each of our pairs is low (Fst less than 0.001 for SD/SP, 0.035 for EF/EA, and 0.071 for France/Egypt). Overall, the consistently strong enrichment of codirectional outliers we observed seems likely to be a product of parallel selective pressures.

To generate hypotheses regarding the genetic basis of parallel adaptation, we performed gene ontology (GO) enrichment analysis for each codirectional comparison. Membrane-associated gene products were enriched in all three comparisons (specifically “membrane part” and “integral to membrane”), and “ion channel activity” was enriched in the EF/FR and FR/SD comparisons. Other categories enriched in two comparisons included “detection of stimulus”, “growth factor activity”, and “innate immune response” (Table S6). Intuitively, the comparison with the strongest codirectional SNP enrichment (EF/FR) had the largest number of GO categories with raw permutation *P* values below 0.05. Enriched terms in this comparison included “neuropeptide receptor activity” (along with other receptor, signaling, and transmembrane terms), “odorant binding”, and “reproduction”, among others (Table S6).

We also performed a search for SNPs with parallel allele frequency change in all three cold-adapted populations. As with the two-pair analyses, an excess of codirectional versus antidirectional *PBE* outliers was observed (Figure 3). Just 28 SNPs had *PBE* quantiles below 0.05 for all population pairs (Table S5), but one of these reached genome-wide significance. This variant showed striking frequency shifts in all three population pairs (Figure 4), with SNP *PBE* quantiles at or below 0.0036 in each case. The random probability of a three-pair codirectional SNP of this type is (1/2)^2^ × (0.0036)^3^ = 1.2 × 10^−8^, such that even with 431,427 tested SNPs, its estimated true positive probability is 99.5% (Table S4). This SNP on chromosome arm 2R is located at position 14,120,310 in release 6 of the *D. melanogaster* genome and position 10,007,815 in release 5. It is located in the first intron of *Prosap*, which encodes a predicted component of the postsynaptic density (Leibl and Featherstone 2008). Curiously, the “cold” allele at this transition polymorphism appears to be the ancestral state, with the same “G” being carried by *D. sechellia, D. simulans, D. erecta*, and *D. yakuba*.

**Figure 4.**
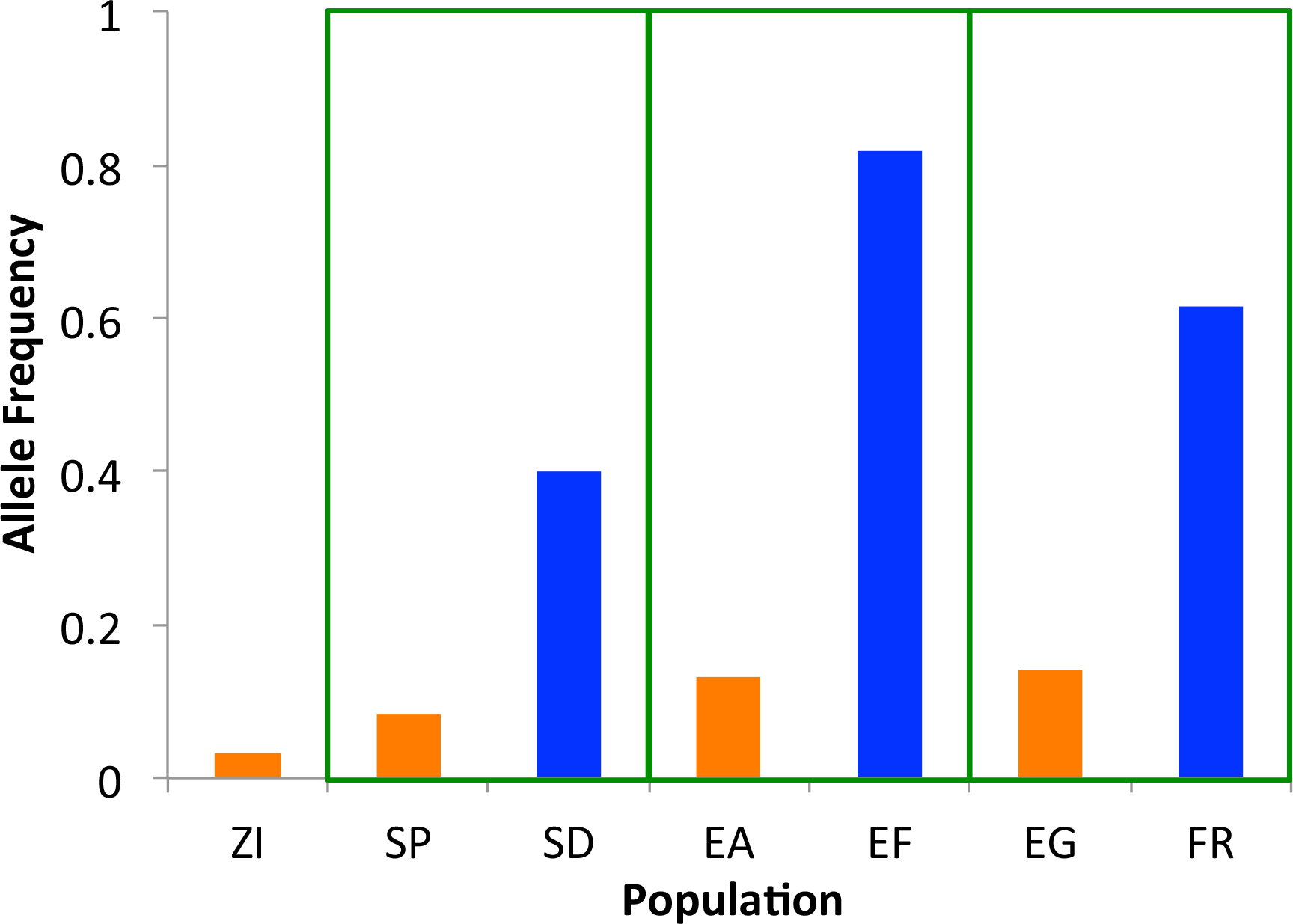
An intronic variant at *Prosap* displays codirectional allele frequency change in three cold-adapted populations (blue bars).

There is little evidence of strong hard sweeps around *Prosap*. The cold-and warm-adapted populations studied all have fairly similar nucleotide diversity in windows across this gene region (Figure S2), including the 4.5 kb window encompassing the codirectional SNP noted above. There are no obvious population differences in haplotype identity (Figure S2), as might be expected after a partial sweep or a moderately soft sweep (Lange and Pool 2016). In terms of window *PBE* values, only the Ethiopian comparison shows some elevation at the focal window (Figure S2). But at this same window, all three comparisons have maximum SNP *PBE* quantiles in the top 5% (Figure S2). This pattern of strong SNP differentiation without a notable linkage signal could reflect soft sweeps (Pennings and Hermisson 2006), which is consistent with the non-trivial frequency of the “cold” allele in the warm-adapted populations.

For France, the parallel divergent SNP highlighted above has the most extreme SNP *PBE* across the entire ~100 kb *Prosap* region (Figure 5). But within a 20 kb region centered on this site, it is only the third and sixth most extreme SNP for South Africa and Ethiopia, respectively. Our alignments also show no evidence for a strongly codifferentiated indel in this region. Hence, although the allele frequency pattern at our focal SNP is dramatic, further study will be needed to confirm which *Prosap* variants may have been under selection in which cold-adapted populations.

**Figure 5.**
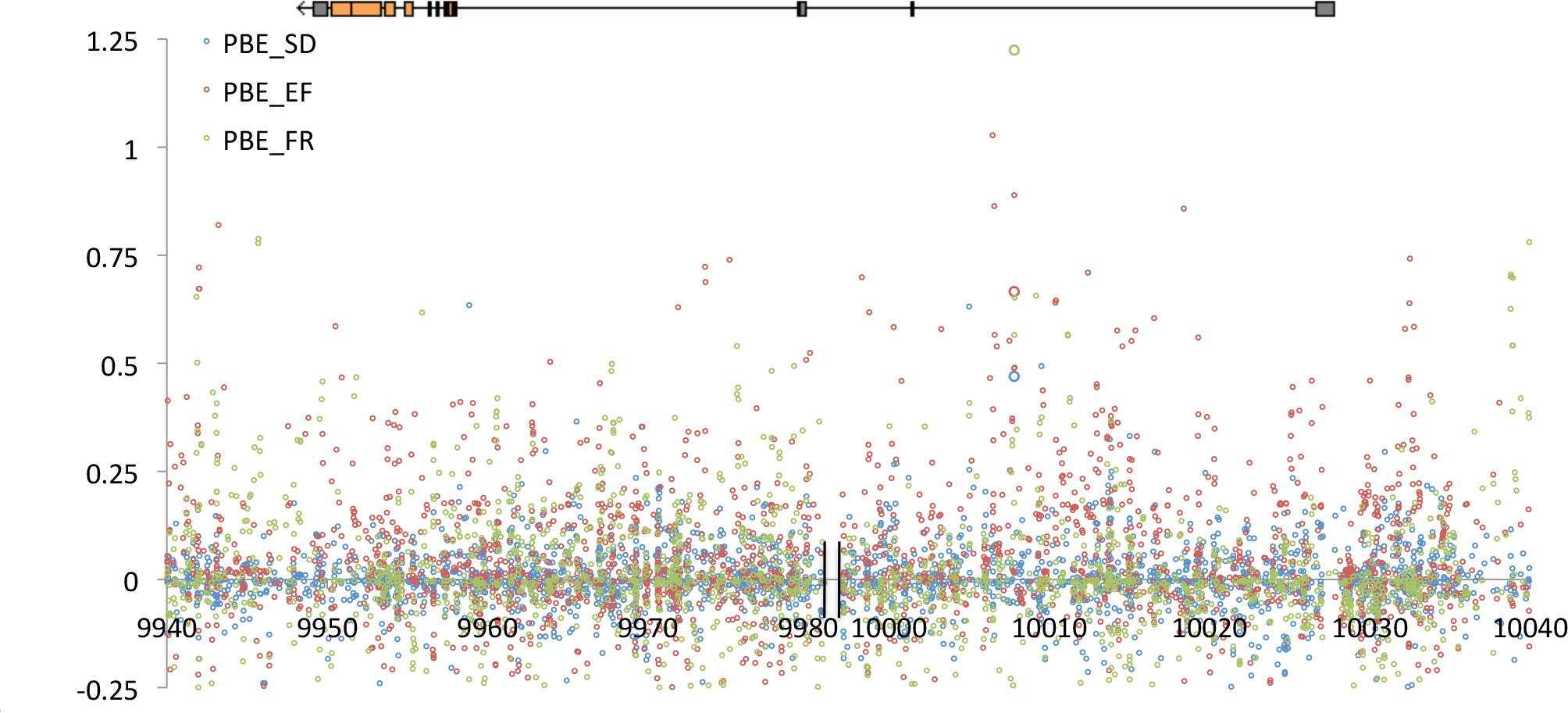
SNP *PBE* across the *Prosap* region is shown for three cold-adapted populations. While our focal SNP (larger circles near 10,008 kb) is the most differentiated for France (green), a handful of other SNPs are more extreme for Ethiopia (red) and South Africa (blue). The top diagram depicts the coding (orange) and non-coding (gray) exons of *Prosap*. The x-axis shows kilobase position along chromosome arm 2R according to release 5 of the *D. melanogaster* reference genome. A break in this axis, 11 kb away from the focal SNP, indicates a 16 kb region in which the reference genome contains two transposable elements (roo and HMS-Beagle) and our reference alignments contain no called sites.

## DISCUSSION

Instances of parallel phenotypic change can shed light on the genetic predictability of evolution. We have shown that as *D. melanogaster* expanded from warm ancestral environments, it evolved greater cold tolerance not only in the palearctic region, but also in the highlands of Ethiopia and South Africa. While natural selection is likely to have favored cold tolerance independently in each case (Introduction), we investigated whether some of the same genetic changes may have facilitated adaptation in two or more colder regions.

We found modest evidence of parallel genetic differentiation in short genomic windows, with an excess of shared outliers specifically between the two African cold-adapted populations. These Ethiopian and South African sample locations are separated by 4,025 km of generally much warmer terrain, so gene flow of cold-adapted alleles between them would require persisting in warm-adapted gene pools. Both of these populations come from high altitude environments (EF at 3,070 meters above sea level, SD at 2,000), so parallel adaptation between them might relate not only to cold, but also to gradients such as air pressure, desiccation, and ultraviolet radiation.

Intriguingly, our SNP-based analysis provided a much more consistent signal of parallel evolution. Overall, SNPs that changed frequency in two different cold-adapted populations (relative to their warm-adapted paired samples) were equally likely to be codirectional (with the same allele rising in each cold-adapted population) or antidirectional. However, SNPs with stronger allele frequency changes were more likely to be codirectional, regardless of the cold-adapted populations compared. It is difficult to exclude all other adaptive and demographic hypotheses, but the clearest explanation for this result would be if cold temperatures or correlated selective pressures favored the same variants in more than one cold-adapted population.

The suggestion that some of the same genetic variants may have been favored in two or more cold-adapted populations raises the question of why our window analysis did not detect a greater number of overlapping *PBE* outliers between cold-adapted populations. The window analysis may reflect a genuine contribution of distinct loci to adaptive evolution in each population. In theory, classic sweep signals could be too narrow (if selection is very weak) or too old for our window analysis to detect. However, divergence among the studied populations may not be ancient enough to facilitate these explanations (Pool *et al.* 2012). Further study of the demographic history of these populations is desirable, although the effects of positive and negative selection on such estimates is not fully clear. Otherwise, the contrasting window and SNP results could reflect an important role for selection on standing genetic variation leading to soft sweeps. If selection favors an existing variant with non-trivial frequency, it may have recombined onto more than one haplotype. The fixation of an allele present on multiple haplotypes has far less influence on linked variation (Pennings and Hermisson 2006). If selection acted on variants with high enough prior frequency, there might be little signal for window *F_ST_* (Lange and Pool 2016) or related statistics like *PBE*. Further study will be needed to identify the evolutionary models and parameters that provide the best match to genetic differentiation at outlier loci across the genome.

There are notable commonalities between the genes and functions identified in our codirectional SNP analysis and in a recent study (Pool 2015). That study examined African and European ancestry along genomes from an admixed population of *D. melanogaster* (the Drosophila Genetic Reference Panel) and found an abundance of “ancestry disequilibrium” in which unlinked loci had correlated ancestry. For example, genomes carrying an African allele at a particular X-linked window might be more likely to carry an African allele in a window on chromosome 2. Of the 17 interaction “hubs” cited there (Figure 5 of Pool 2015), 10 of the listed genes were also detected in at least one of our codirectional comparisons (*AstA-R1, CcapR, CCHa1-R, Fife, GABA-B-R1, norpA, Or65b, Piezo, Rdl, shakB*, and *Spn*). These genes include several with roles in neurotransmission, which is also influenced by the postsynaptic density to which *Prosap* contributes (Liebl and Featherstone 2008). The above genes were previously identified not only by ancestry disequilibrium, but also by elevated *F_ST_* between European and western African genomes (Pool 2015), so their detection in this study is not entirely independent. However, our results raise the possibility that similar genes and processes may have been under selection in other cold-adapted populations as well.

Prior to this study, GO categories related to the nervous system have been enriched in a variety of genome scans for positive selection in *D. melanogaster* (Langley *et al.* 2012; Pool *et al.* 2012; Pool 2015). And yet, very few empirical studies have investigated nervous system evolution in *Drosophila*. A rare example is the study of Campbell and Ganetzky (2012), who found that the structure of a neuromuscular junction (a synapse between a neuron and a muscle fiber) has evolved dramatically between *Drosophila* species, even when closely related taxa were compared. We hypothesize that parallel nervous system evolution may have occurred within *D. melanogaster* during adaptation to cold temperatures or a correlated selective pressure. Such evolution may have served to maintain ancestral function in a challenging environment, or it may have adaptively altered behavior or other phenotypes. Expanded study of nervous system evolution in *D. melanogaster* is needed to clarify the molecular mechanisms and phenotypic relevance of parallel adaptive changes at these genes.

## MATERIALS AND METHODS

### Fly strains and phenotypic assays

All fly population samples used here were described in previous population genomic studies (Pool *et al.* 2012; Lack *et al.* 2015). The strains used in this study were derived from independent isofemale lines and inbred for eight generations prior to study. Cold tolerance experiments focused on females that were 4 to 7 days old at the outset of the experiment. Prior to cold exposure, flies were raised at 25°C on medium prepared in batches of: 4.5 L water, 500 mL cornmeal, 500 mL molasses, 200 mL yeast, 54 g agar, 20 mL propionic acid, and 45 mL tegosept 10% (in 95% ethanol). To control larval density, flies were raised in half pint bottles with 20 virgin females and 20 males allowed to oviposit for 48 hours.

The cold tolerance experiment involved placing flies in a 4°C chamber. For each strain, a bottle containing 30-50 females was used. After 96 hours, the number of standing and knocked-down flies in each glass vial was counted. Flies remaining on their legs were deemed to retain mobility, while other flies were considered immobile (dead or in chill coma). In our hands, we found that this 4°C assay provided more repeatable results than chill coma response time or survival after brief exposure to below-freezing temperatures.

Most testing was done on inbred lines. For a subset of lines from each population sample, we investigated cold tolerance in outbred flies as well. For each population sample, a single inbred line was used as the source of virgin females for between-line crosses. This “reference line” was then crossed to males from a series of other inbred lines from the same population sample. The cold tolerance observed for outbred F1 flies from each of these crosses was compared to expectations based on the midparent value from the two parental inbred strains.

### Genomes and inversion testing

An initial set of genomes from each of these six population samples was previously published (Pool *et al.* 2012; Lack *et al.* 2015). Additional genomes used here (from a subset of previously studied populations) were sequenced, aligned, and filtered (for heterozygosity, identity by descent, and population admixture) using identical methods to those described previously (Lack *et al.* 2015). These genomes are documented in an accompanying article that expands the *Drosophila* Genome Nexus (Lack *et al.* 2016a). Recent gene flow into Africa, which may reflect urban adaptation, was masked from this data set (Pool *et al.* 2012; Lack *et al.* 2015; Lack *et al.* 2016a).

Our study focuses on three pairs of population samples, in which the closely-related members of each pair derive from warm and cold environments, respectively. The population genetic analyses described below also draw upon a published sample of genomes from Zambia (ZI), which is thought to represent an ancestral range population (Pool *et al.* 2012; Lack *et al.* 2015). Numbers of genomes analyzed for each population are given in Table 1.

We used PCR to test inbred strains from each population sample for eight common inversions (*In1A, In1Be, In2Lt, In2RNS, In3LP, In3RK, In3RMo*, and *In3RP*), using primers and conditions given by Lack *et al.* (2016b). These primers were modified from those published by Corbett-Detig *et al.* (2012) to accommodate or avoid common SNPs.

### Window-based population genetic analysis

Our primary interest in genomic analysis is to identify loci with strong genetic differentiation between the members of a warm-and cold-adapted population pair, which may represent targets of population-specific positive selection. The primary population genetic approach uses statistics based on *F_ST_*, which quantifies allele frequency differences between two populations. By adding pairwise *F_ST_* values involving a third “outgroup” population, *PBS* isolates the allele frequency change that is specific to one focal population’s history (Yi *et al.* 2010). *PBS* will be large when natural selection has been specific to the focal population, but it will also be large when all populations have long branches. In this study, our interest is to find loci that experienced positive selection in the cold-adapted populations specifically, and we would like to avoid universally-selected loci when testing whether different cold-adapted populations share many population genetic outliers in common. We therefore used a recently-described variant of *PBS* known as Population Branch Excess (*PBE;* Yassin *et al.* 2016). *PBE* quantifies the difference between the observed *PBS* value and that expected based on (1) genetic differentiation between the two non-focal populations at this locus, and (2) typical values for that quantity and *PBS* at other loci. Numerically,

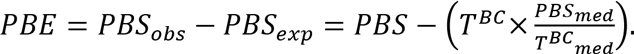

Here, *T^BC^* quantifies genetic differentiation between the non-focal populations, as −log(1 − *F_ST_*), which scales our locus-specific expectations for genetic differentiation. The ratio of median values reflects the typical relationship between *PBS* and *T^BC^* observed genome-wide (or in our case, for a given chromosome arm). We expect *PBE* to be strongly positive when selection has been specific to the focal population, but not when selection has increased all branch lengths similarly.

We calculated *PBE* for genomic windows with sizes scaled by local diversity levels. Each window contained 250 non-singleton single nucleotide polymorphisms (SNPs) in the ZI population. These windows had a median size of roughly four kilobases. To focus on regions in which selection signals should be more localized, we focused our population genetic analyses on middle portions of chromosome arms where sex-averaged crossing-over rates are generally above 0.5 cM / Mb (Comeron *et al.* 2012), specifically: 2.3 to 21.4 Mb for the X chromosome, 0.5 to 17.5 Mb for arm 2L, 5.2 to 20.8 Mb for arm 2R, 0.6 to 17.7 Mb for arm 3L, and 6.9 to 26.6 Mb for arm 3R. We also evaluated *PBE* for specific SNPs, and identified the maximum SNP *PBE* value for each window. For both window and maximum SNP *PBE*, we computed quantiles corresponding to the proportion of analyzed windows on the same chromosome arm with an equal or greater value of *PBE*. All window and SNP-specific analyses included only genomes with a standard arrangement inferred for the chromosomal arm of interest (Corbett-Detig *et al.* 2012).

We used a shift permutation approach to test whether window *PBE* outliers in different populations overlapped more than expected. Outlier windows were grouped into “regions” encompassing all neighboring windows with *PBE* quantile below 0.05, interrupted by no more than four non-outlier windows. Comparing two sets of outlier regions, one set was shifted within chromosome arms. Outlier region locations were shifted in increments of five windows to increase the independence of shift replicates. Outliers were shifted a minimum of 50 windows from their empirical locations. Based on the windows present on the shortest chromosome arm (2R), this allowed for a total of 827 shift permutations.

### SNP-based population genetic analysis

Analogous to the study of Tennessen and Akey (2011), we tested whether SNP frequency differences tend to have the same allele at elevated frequency in two or more cold-adapted populations. Specifically, we assigned a SNP quantile to the *PBE* value for each of a population pair’s analyzed SNPs. When then tested whether “codirectional” SNPs (those with the same allele at elevated frequency in both cold-adapted populations relative to their warm-adapted counterparts) were more common than “antidirectional” SNPs in which opposite alleles were at higher frequency in the two cold-adapted populations. Non-directional SNPs (those with no frequency change in one or both pairs) were excluded from this analysis, so the null expectation is that 50% of SNPs should be codirectional by chance. If parallel selective pressures have raised the frequency of certain variants in more than one cold-adapted population, we might observe that SNPs with high *PBE* in both population pairs are more likely to be codirectional than antidirectional. Here, we focused on the comparison of SNPs with *PBE* quantile below 0.05 in both population pairs versus those with *PBE* quantile above 0.1 in one or both pairs. We also performed a codirectional analysis with all three population pairs, using the same data filters outlined above. Here, the proportion of codirectional SNPs is expected to be one quarter by chance, rather than one half.

Within each pair of cold-and warm-adapted populations, we excluded sites with less than ten called alleles in the cold-adapted population, or less than five called alleles in each warm-adapted population. We restricted the analysis to SNPs with an average population frequency of the minor allele of at least 10% between the two populations. We then created a merged set of SNPs that met our sampling and frequency criteria for each of two population pairs. Although SNPs with intermediate *versus* rare frequency in the ancestral population might have different distributions of *PBE* under neutrality, we would still expect both classes to be codirectional and antidirectional with equal probability.

We performed gene ontology enrichment analysis on SNPs identified as codirectional outliers, separately for each of the three analyses comparing two population pairs (EF/FR, EF/SD, and FR/SD with respect to the cold-adapted populations). Outlier SNP lists were pared to remove sites within 10 kb of each other, excluding the site with the least extreme *PBE* quantile in either population pair – in other words, retaining the site with the lower max(*PBEQ1, PBEQ2*). For exonic SNPs (coding or UTR), the genes with overlapping exons were noted. For non-exonic SNPs, the genes with the nearest proximal and distal exons were noted. Each outlier SNP could only implicate a specific GO category once, and each gene could only be counted once. Permutations (50,000) were performed in which each outlier SNP on the pared list was randomly relocated within the analyzed region of the same chromosome arm (thus accounting for variability in gene length). For each GO category, a *P* value indicated the proportion of permutation replicates in which an equal or greater number of genes was implicated.

## ACKNOWLEDGMENTS

This work was funded by the National Institutes of Health through a grant to JEP [R01 GM111797] and a fellowship to JBL [F32 GM106594].

## REFERENCES

Ayrinhac A, Debat V, Gibert P, Kister A-G, Legout H, Moreteau B, Vergilino R, David JR. 2004. Cold adaptation in geographical populations of Drosophila melanogaster. phenotypic plasticity is more important than genetic variability. Func Ecol. 18.700–706.

Bergland AO, Tobler R, Gonzalez J, Schmidt P, Petrov D. 2016. Secondary contact and local adaptation contribute to genome-wide patterns of clinal variation in Drosophila melanogaster. Mol Ecol. 25.1157–1174.

Božičević V, Hutter S, Stephan W. 2016. Population genetic evidence for cold adaptation in European Drosophila melanogaster populations. Mol Ecol. 25.1175–1191.

Campbell M, Ganetzky B. 2012. Extensive morphological divergence and rapid evolution of the larval neuromuscular junction in Drosophila. Proc Natl Acad Sci USA. 109.E648–E655.

Cavalli-Sforza L. 1969. Human diversity. Proc 12th Int Congr Genet. 2.405–416.

Comeron JM, Ratnappan R, Bailin S. 2012. The many landscapes of recombination in Drosophila melanogaster. PLoS Genet 8.e1002905.

Corbett-Detig RB, Cardeno C, Langley CH. 2012. Sequence-based detection and breakpoint assembly of polymorphic inversions. Genetics 192.131–137.

Dahlgaard J, Hoffmann AA. 2000. Stress resistance and environmental dependency of inbreeding depression in Drosophila melanogaster. Cons Biol. 14.1187–1192.

DePristo M, Banks E, Poplin R, Garimella K, Maguire J, Hartl C, Philippakis AA, Del Angel G, Rivas MA, Hanna M, et al., 2011. A framework for variant discovery and genotyping using next-generation DNA sequencing data. Nature Genet. 43. 491–498.

Dieringer D, Nolte V, Schlötterer C. 2005. Population structure in African Drosophila melanogaster revealed by microsatellite analysis. Mol Ecol. 14.563–573.

Duchen P, Živković D, Hutter S, Stephan W, Laurent S. 2013. Demographic inference reveals African and European admixture in the North American Drosophila melanogaster population. Genetics. 193:291–301.

Hancock AM, Witonsky DB, Ehler E, Alkorta-Aranburu G, Beall C, Gebremedhin A, Sukernik R, Utermann G, Pritchard J, Coop G, et al. 2010. Human adaptations to diet, subsistence, and ecoregion are due to subtle shifts in allele frequency. Proc Natl Acad Sci USA 107:8924–8930.

Hoffmann AA, Sørensen JG, Loeschcke V. 2003. Adaptation of Drosophila to temperature extremes: bringing together quantitative and molecular approaches. J Therm Biol. 28:175–216.

Kao JY, Zubair A, Salomon MP, Nuzhdin SV, Campo D. 2015. Population genomic analysis uncovers African and European admixture in Drosophila melanogaster populations from the south-eastern United States and Caribbean Islands. Mol Ecol. 24:1499–1509.

Kapun M, Fabian DK, Goudet J, Flatt T. 2016. Genomic evidence for adaptive inversion clines in Drosophila melanogaster. Mol Biol Evol. 33:1317–1336.

Kumar S, Stecher G, Tamura K. 2016. MEGA7: Molecular Evolutionary Genetics Analysis version 7.0 for bigger datasets. Mol Biol Evol. In Press.

Lachaise D, Cariou M-L, David JR, Lemeunier F, Tsacas L, Ashburner M. 1988. Historical Biogeography of the Drosophila melanogaster Species Subgroup. In: Hecht MK, Wallace B, Prance GT, editors. Evolutionary Biology. Boston, MA: Springer. p. 159–225.

Lack JL, Cardeno CM, Crepeau MW, Taylor W, Corbett-Detig RB, Stevens KA, Langley CH, Pool JE. 2015. The Drosophila genome nexus: a population genomic resource of 623 Drosophila melanogaster genomes, including 197 from a single ancestral range population. Genetics. 199:1229–1241.

Lack JL, Lange JD, Tang AD, Corbett-Detig RB, Pool JE. 2016. A thousand fly genomes: version 1.1 of the Drosophila Genome Nexus. Mol Biol Evol., co-submitted with this article.

Lack JB, Monette MJ, Johanning EJ, Sprengelmeyer QD, Pool JE. 2016. Decanalization of wing development accompanied the evolution of large wings in high-altitude Drosophila. Proc Natl Acad Sci USA 113:1014–1019.

Lange JD, Pool JE. 2016. A haplotype method detects diverse signatures of local adaptation from genomic sequence variation. Mol Ecol. In Press.

Lemeunier F, Aulard S. 1992. Inversion polymorphism in Drosophila melanogasters. In: Krimbas CB, Powell JR, editors. Drosophila inversion polymorphism. Boca Raton (FL): CRC Press. p. 339–405.

Li H, Durbin R. 2010. Fast and accurate long-read alignment with Burrows-Wheeler transform. Bioinformatics 26:589–595.

Liebl FLW, Featherstone DE. 2008. Identification and investigation of Drosophila postsynaptic density homologs. Bioinform Biol Insights 2:369–381.

Lunter G, Goodson M. 2010. Stampy: a statistical algorithm for sensitive and fast mapping of Illumina sequence reads. Genome Res. 18:821–829.

Ohta T. 1971. Associative overdominance caused by linked detrimental mutations. Genet Res. 18:277–286.

Pennings PS, Hermisson J. 2006. Soft sweeps III: the signature of positive selection from recurrent mutation. PLoS Genet. 2:e186.

Pool JE. 2015. The mosaic ancestry of the Drosophila Genetic Reference Panel and the D. melanogaster reference genome reveals a network of epistatic fitness interactions. Mol Biol Evol. 32.3236–3251.

Pool JE, Corbett-Detig RB, Sugino RP, Stevens KA, Cardeno CM, Crepeau MW, Duchen P, Emerson JJ, Saelao P, Begun DJ, et al. 2012. Population genomics of sub-Saharan Drosophila melanogaster. African diversity and non-African admixture. PLoS Genet. 8.e1003080.

Svetec N, Werzner A, Wilches R, Pavlidis P, Alvarez-Castro JM, Broman KW, Metzler D, Stephan W. 2011. Identification of X-linked quantitative trait loci affecting cold tolerance in Drosophila melanogaster and fine mapping by selective sweep analysis. Mol Ecol. 20.530–544.

Thornton KR, Andolfatto P. 2006. Approximate Bayesian inference reveals evidence for a recent, severe bottleneck. in a Netherlands population of Drosophila melanogaster. Genetics. 172.1607–1619.

Turner TL, Levine MT, Eckert ML, Begun DJ. 2008. Genomic analysis of adaptive differentiation in Drosophila melanogaster. Genetics. 179.455–473.

Yassin A, Debat V, Bastide H, Gadesweski N, David JR, Pool JE. 2016. Repeated ecological specialization and incipient speciation in an island Drosophila population. Proc Natl Acad Sci USA 113.4771–4776.

Yi X, Liang Y, Huerta-Sanchez E, Jin X, Cuo ZXP, Pool JE, Xu X, Jiang H, Vinckenbosch N, Korneliussen TS, et al. 2010. Sequencing of 50 human exomes reveals adaptation to high altitude. Science. 329.75–78.

